# Perinatal Nicotine Exposure Disrupts Hematopoietic Stem Cell Development and Elevates Influenza Susceptibility in Adulthood

**DOI:** 10.1101/2025.02.23.639728

**Authors:** T Cool, A Rodriguez y Baena, MGE Rommel, C Mattingly, E Bachinsky, S Saini, S Chattopadhyaya, BA Manso, S Rajendiran, AK Worthington, DM Poscablo, A Deguzman, T Berger-Cahn, DF Boyd, EC Forsberg

## Abstract

Tobacco use during pregnancy has many deleterious health consequences for not only the smoking mother, but also on the unborn fetus. Children of smoking mothers are reported to have higher frequency and severity of respiratory diseases later in life; however, the mechanisms driving this increased vulnerability are not clearly understood. One potential cause of increased disease susceptibility is an altered immune system, originating in epigenetically maladaptive hematopoietic stem cells (HSCs). Here, we show that perinatal nicotine exposure (PNE) alters the establishment of HSCs and fetal-derived non-traditional tissue immune cells, with no alterations in circulating immune cell numbers. Suppression of HSCs and lung immune cells persisted for weeks after PNE had ceased. Strikingly, PNE led to increased disease susceptibility and severity upon challenge with influenza A virus in adulthood. This was associated with significant and highly selective alterations in lung immune cells, emphasizing the importance of cellular mechanisms in resilience to infections. Together, these experiments demonstrate that perinatal exposures that have deleterious consequences on hematopoietic establishment can impair immune function for life and identify the cellular mechanisms by which perinatal nicotine exposure predisposes the offspring to a weakened defense against respiratory pathogens.

**HIGHLIGHTS:**

- Perinatal nicotine exposure (PNE) causes long-term alterations of hematopoiesis
- PNE perturbs hematopoietic stem cell (HSC) development and maintenance
- PNE diminishes the population of fetal-derived tissue-resident alveolar macrophages
- PNE alters cellular mechanisms, exacerbating disease severity later in life
- Nicotine suppression of central immunity is manifested mechanistically by altered immune cell output

## INTRODUCTION

The immune system is established through developmental immune layering, a complex orchestration of unique waves of hematopoietic stem and progenitor cells (HSPCs) that give rise to developmentally distinct subsets of immune cells (Cool & Forsberg, 2019). This process comprises non-self-renewing progenitors that generate long-lived self-renewing mature cells, such as tissue-resident macrophages (Cool & Forsberg, 2019; Guilliams et al., 2013; Lichanska & Hume, 2000; Sorokin et al., 1992). Several perinatally-established subsets of immune cells, including tissue-resident macrophages and innate-like lymphoid cells, have been implicated as modulators of inflammation across tissues (Hildreth & O’Sullivan, 2019; Lazarov et al., 2023; Vivier et al., 2018), but the extent to which these fetal-derived immune cells and their progenitor source persist and contribute to adult immunity remains unclear. Starting at mid-gestation (∼E10.5), self-renewing progenitors, such as traditional hematopoietic stem cells (HSCs), give rise to non-self-renewing progeny which are replenished and sustained from adult bone marrow (BM) HSCs for life (Boisset et al., 2010; Cool & Forsberg, 2019; Lewis et al., 2021). The recent concept of “central trained immunity” has demonstrated that HSCs retain an epigenetically established immune memory that shapes long-term immune cell production (de Laval et al., 2020; Divangahi et al., 2021; Johansson et al., 2023; Kain et al., 2023; Kaufmann et al., 2018; Netea et al., 2020). The timing of in utero exposure to pathogens and toxicants coincides with HSC establishment and with distinct developmental waves that generate long-lasting immune cells. Therefore, self-renewing HSCs and long-lived immune cells may serve as key mediators between early-life environmental exposures and increased disease susceptibility later in life.

Accumulating evidence has shown that perinatal exposure to environmental toxins, including tobacco smoke, affects perinatal immune establishment and function leading to long-lasting consequences (Avşar et al., 2021; J. Cao et al., 2016; Cool et al., 2022; Dietz et al., 2010; Karthikeyan et al., 2024; Quelhas et al., 2018). Nicotine, the major component of tobacco products, is a nicotinic receptor agonist and its binding triggers cellular responses such as proliferation, apoptosis, and differentiation (Ben-Yehudah et al., 2013; Cool et al., 2022; Piao et al., 2009; Schaal & Chellappan, 2014). Additionally, nicotine is known to cause inflammation (Gracia, 2005; Mohsenzadeh et al., 2014) which can lead to permanent alterations in immunity later in life (Apostol et al., 2020; López et al., 2023; Pattenden et al., 2006). However, the cellular and molecular mechanisms underlying the increased susceptibility to respiratory disease among children of smoking mothers are poorly understood. Here, we implemented an *in vivo* model to investigate how perinatal nicotine exposure (PNE) alters hematopoietic establishment and life-long function in the adult offspring.

## RESULTS

### Perinatal nicotine exposure alters seeding of HSCs in the fetal liver

To determine how PNE affects hematopoiesis, we administered nicotine (100 µg/mL) in sucrose solution *ad libitum* via drinking water (Chang et al., 2010; A. C. Collins et al., 2012; Isotani et al., 2025; Maier et al., 2011) to pregnant dams for the entirety of gestation (**Figure 1A**). There were no differences in body weight at P0 between control and nicotine-exposed pups (**Supplementary Figure 1B**). Then, we analyzed the offspring at postnatal day 0 (P0) to determine whether PNE affected hematopoiesis in the two major hematopoietic compartments at this timepoint, which are the liver and the bone marrow (BM) (A. Collins et al., 2024). We observed a decrease in the total number of cells in the liver (**Supplementary Figure 1C**), and, as a potential cause of this decrease, that the number of HSCs was significantly decreased in the liver of nicotine-exposed P0 pups compared to control pups (**Figure 1B, Supplementary Figure 1A**). In contrast, the number of HSCs seeded in the BM was significantly increased in nicotine-exposed pups compared to controls (**Figure 1C**). This increase in BM HSC numbers did not fully account for the decreased HSC numbers in the liver of these pups as the total HSC number in the P0 pups was still significantly reduced in the nicotine group (**Supplementary Figure 1D**). Moreover, the cell cycle profile of liver HSCs (**Figure 1D**) and BM HSCs (**Supplementary Figure 1E**) remained unaltered, suggesting that HSC proliferative exhaustion is an unlikely reason for reduced HSC numbers at this time point. To test whether the observed decrease in HSC number at P0 was due to direct cytotoxic effects following nicotine binding to one of its receptors, we quantified the expression of the nicotinic receptor α7 (nAChRα7), which was previously reported to be expressed on hematopoietic cells (Chang et al., 2010). We did not detect any transcriptional expression of nAChRα7 in HSCs from either control or nicotine-exposed P0 pups. Instead, we detected receptor expression in whole liver tissue from both from control and nicotine-exposed P0 pups (**Supplementary Figure 1F**), suggesting that nicotine might interact with liver niche cells instead of directly affecting HSCs. Thus, we investigated whether PNE affected the liver microenvironment. We collected serum (extracellular fluid) from the liver of P0 pups and measured an array of cytokines and chemokines secreted within the liver niche by multiplex ELISA (**Figure 1E**). These data were normalized to the total protein concentration of each liver (**Supplementary Figure 1G**). While the levels of some pro-inflammatory cytokines such as IL-1β, IL-6, and IFNβ remained unaffected (**Figure 1F**), other molecules previously shown to be affected by nicotine or cigarette smoke followed similar patterns in this screen, such as MIG (CXCL9) (Wang et al., 2022) being significantly higher (**Figure 1G**) and TIMP-1 (Katono et al., 2006; Watson et al., 2010), TNFα (Kizildag et al., 2021; Li et al., 2011), and MCP-1 (Valdez-Miramontes et al., 2020) being significantly lower (**Figure 1H**) in nicotine-exposed pups compared to controls. These changes in cytokine and chemokine expression suggest that PNE can shape the fetal liver microenvironment, potentially affecting the composition of the liver and/or altering migration, proliferation, and extravasation of immune cells typically present in the fetal liver. This was supported by the increased liver weights in nicotine-exposed pups (**Supplementary Figure 1H**) (Konno et al., 2020). These data suggest that nicotine-induced changes by cytokines and chemokines in the perinatal liver microenvironment can alter the HSC pool, potentially leading to lasting alterations of the hematopoietic system.

**Figure 1.**
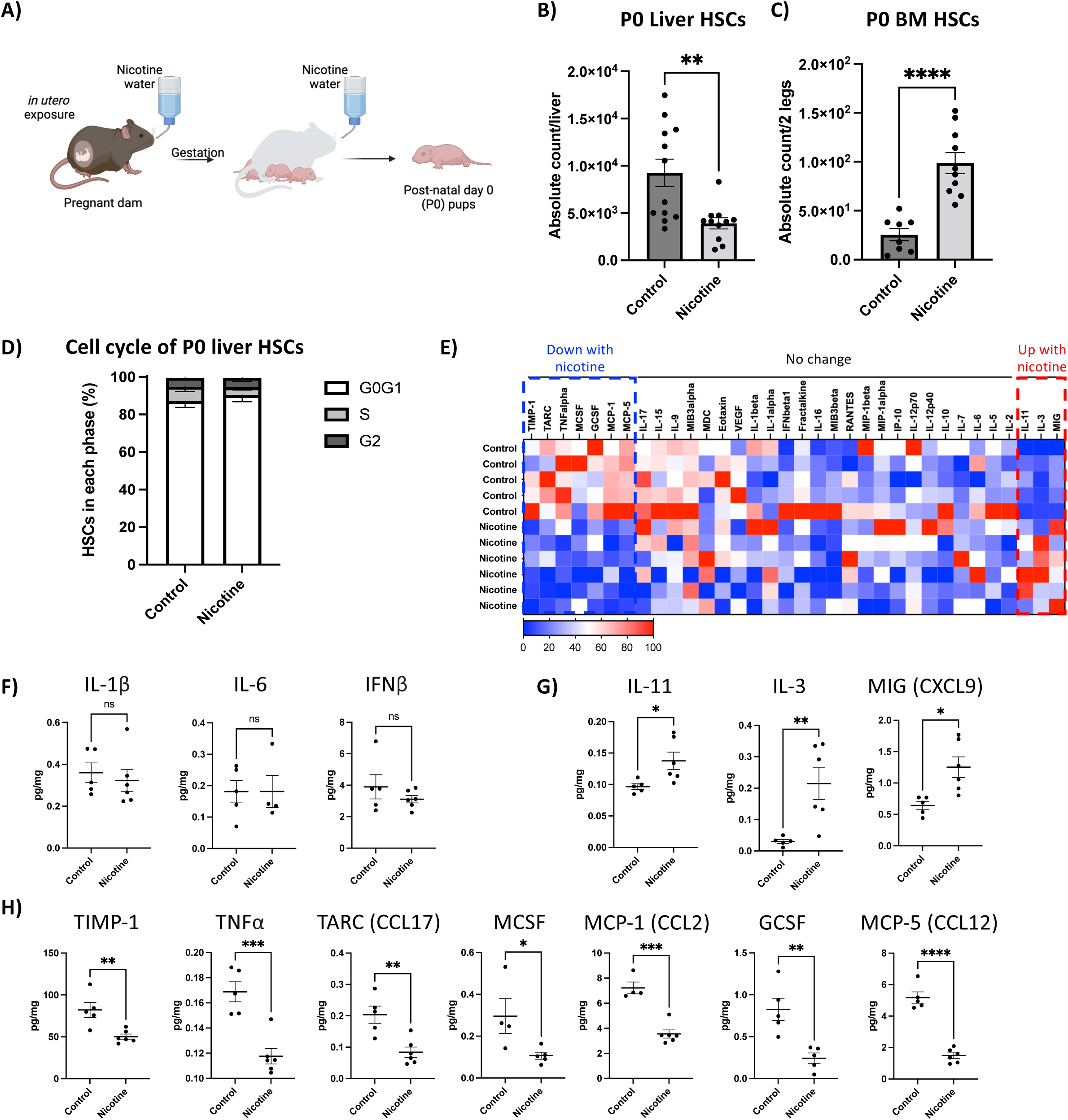
Perinatal nicotine exposure (PNE) alters HSC number and liver niche in newborn pups. A) Experimental set up of PNE. B) PNE decreases the numbers of HSCs in the fetal liver. Absolute cell counts of liver HSCs from control (dark gray bars, n=12) or nicotine-exposed (light gray bars, n=11) post-natal day 0 (P0) pups; **P<0.01 (Student’s t-test). C) PNE increases the numbers of HSCs in the fetal bone marrow. Absolute cell counts of bone marrow (BM) HSCs from control (n=8) or nicotine-exposed (n=10) P0 pups. ****P<0.001 (Student’s t-test). D) PNE does not significantly alter the cell cycle status of fetal liver HSCs. Cell cycle status of liver HSCs from control (n=6) or nicotine-exposed (n=6) P0 pups E-H) PNE selectively alters the fetal liver cytokine environment. E) Heat-map representation of concentration for 35 cytokines/chemokines arrayed in liver serum from control (n=5) or nicotine-exposed (n=6) P0 pups. Concentrations were scaled and normalized within groups for each cytokine (shown as blue=low and red=high); the actual concentrations (pg/mg) are shown in F-H. F-H) Concentrations of cytokines/chemokines that were unchanged (F), significantly higher (G), or significantly lower (H) in liver serum from nicotine-exposed pups compared to control. ns, not significant; *P<0.05, **P<0.01, ***P<0.005, ****P<0.001 (Student’s t-test). Data are mean+/-SEM; each dot represents an individual animal.

### Perinatal nicotine exposure permanently reduces numbers of HSPCs in the bone marrow

To determine if changes within the HSC compartment of P0 pups exposed to nicotine in utero (**Figure 1**) persist until later time points, we analyzed the offspring following PNE at post-natal day 14 (P14) and at 8-12 weeks (young adulthood) (**Figure 2A**). We analyzed the BM as it is the main hematopoietic organ at these time points (A. Collins et al., 2024). We observed no differences in body weight of control and nicotine-exposed P14 pups (**Figure 2B**), suggesting that the offspring were feeding and developing properly. Interestingly, at P14, the number of BM HSCs was significantly lower in the nicotine-exposed pups (**Figure 2C**), reversing the increase observed at P0 (**Figure 1C**) and instead reflecting the overall decrease in HSCs when accounting for fetal liver HSCs (**Figure 1B and Supplementary Figure 1D**). This decrease in HSC numbers was reflected in other hematopoietic progenitor populations, namely multipotent progenitors (MPPs: Lin-, ckit+, Sca1+, CD150-, Flk2+) (**Figure 2D**) and myeloid progenitors (MyPros: Lin-, cKit+, Sca1-) (**Figure 2E**), which can be phenotypically subdivided into common myeloid progenitors (CMPs: Lin-,cKit+, Sca1-, CD16/32mid, CD34+), granulocyte-monocyte progenitors (GMPs: cKit+, Sca1-, CD16/32+, CD34+), and megakaryocytic-erythroid progenitors (MEPs: cKit+, Sca1-, CD16/32-, CD34-) (**Supplementary Figure 2B-D**). The reduction in HSPC numbers was not reflected in an overall decrease in immature (lineage-negative) cells in the BM of nicotine-exposed pups (**Supplementary Figure 2A**). Additionally, similar to the P0 time point, we did not observe any changes in cell cycling of HSCs at P14 (**Figure 2F**). We then analyzed the BM composition of adult offspring that had been exposed to nicotine perinatally. Interestingly, we observed that at this time point the number of HSCs remained significantly lower in the nicotine-exposed mice compared to control (**Figure 2G**). In contrast, we observed no significant difference in the number of live, lineage negative BM cells (**Supplementary Figure 2E**), MPPs (**Figure 2H**) or MyPros (**Figure 2I**; **Supplementary Figure 2F-H**) between control and experimental mice. No difference in the cell cycle status of adult HSCs was observed (**Figure 2J**). To determine whether the developmental age during which mice are exposed to nicotine differentially affects hematopoiesis, we exposed adult (8-12 weeks) mice to nicotine (100 µg/mL) in sucrose solution *ad libitum* via drinking water for ∼8 weeks (**Figure 2K**; these mice had not been exposed to nicotine perinatally). The effect of adult exposure on HSCs was not concordant with what we observed by PNE. Instead, we observed a significant increase in the number of phenotypic HSCs in the BM of nicotine-treated adult mice (**Figure 2L**) along with a significant increase in white blood cell (WBC) counts (**Figure 2O**). These results are consistent with previous reports (Chang et al., 2010; Zalokar et al., 1981). Meanwhile, adult-exposed MPPs and MyPros remained unaffected by nicotine treatment (**Figure 2M-N**). Taken together, these data suggest that the developmental age during which mice are exposed to nicotine differentially affects HSCs and hematopoiesis. Specifically, there was a long-lasting decrease of BM HSC and HSPC numbers upon perinatal nicotine exposure, in stark contrast with the selective increase in numbers of HSCs observed upon nicotine exposure in adulthood.

**Figure 2.**
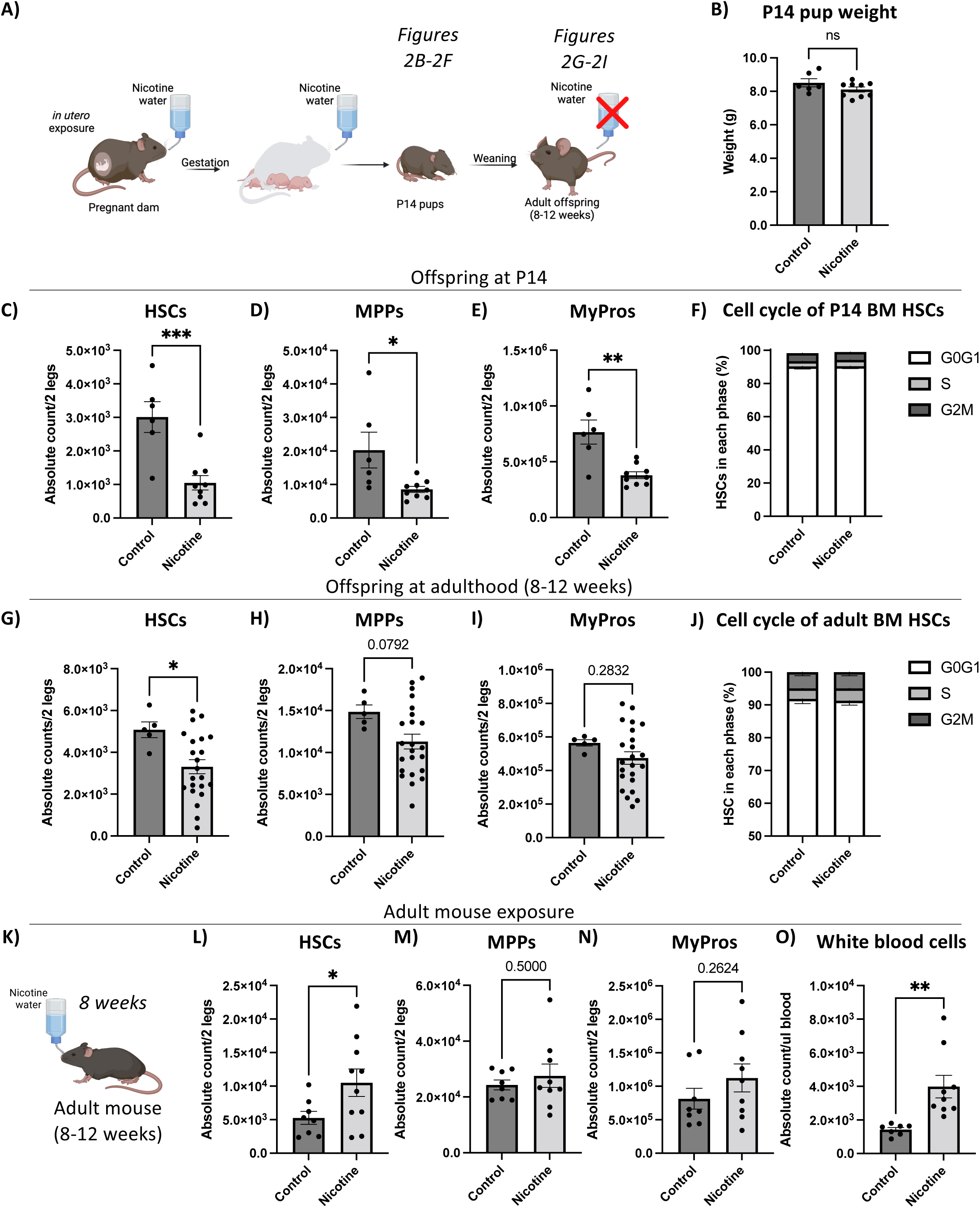
Perinatal nicotine exposure permanently reduced BM HSC numbers. A) Experimental set up of PNE. B) Body weight of P14 pups from control (n=6) and nicotine-exposed groups (n=9). C-E) PNE reduces the numbers of hematopoietic stem and progenitor cells in P14 BM. Absolute cell counts of BM HSCs (C), MPPs (D), and MyPros (E) from the same mice as (B). *P<0.05, **P<0.01, ***P<0.005 (Student’s t-test). F) Cell cycle status of P14 BM HSCs from control (n=6) or nicotine-exposed (n=6) P14 pups. C-E) G-I) PNE selectively reduces the adult HSC numbers. Absolute cell counts of BM HSCs (G), MPPs (H), and MyPros (I) from control (n=5) or nicotine-exposed (n=24) adult offspring. *P<0.05 (Student’s t-test). J) PNE does not significantly alter the cell cycle status of adult HSCs. Cell cycle status of adult (8-12 weeks) BM HSCs from control (n=6) or nicotine-exposed (n=7) mice. K) Experimental set up of adult nicotine exposure. L-O) Adult nicotine exposure selectively increases the numbers of BM HSCs and circulating white blood cells. Absolute cell counts of bone marrow HSCs (L), MPPs (M), MyPros (N), and peripheral blood white blood cells (O) from control (n=8) and nicotine-exposed (n=10) adult mice. *P<0.05, **P<0.01 (Student’s t-test). Data mean +/- errors bars represent SEM; each dot represents an individual animal.

### Perinatal nicotine exposure does not affect mature cell numbers in the peripheral blood

To determine if the changes observed within the BM HSPC compartment were reflected in circulating cell populations, we analyzed the peripheral blood of P14 and 8-12 week old adult offspring that had been exposed to nicotine *in utero* (**Figure 2A**). Interestingly, although PNE led to altered BM HSPC numbers, we observed no significant differences in the number of myeloid cells (“GMs”, CD11b+Gr1+ cells; **Figure 3A**), B cells (**Figure 3B**), T cells (**Figure 3C**), red blood cells (RBCs) (**Figure 3D**), or platelets (**Figure 3E**) of P14 pups. Similarly, we observed no differences in cell numbers in the peripheral blood of adult offspring of nicotine-exposed mothers (**Figure 3F-J**). Thus, the decreased numbers within the BM HSPC compartment of nicotine-exposed offspring were able to sustain mature blood cell homeostasis in the peripheral blood.

**Figure 3.**
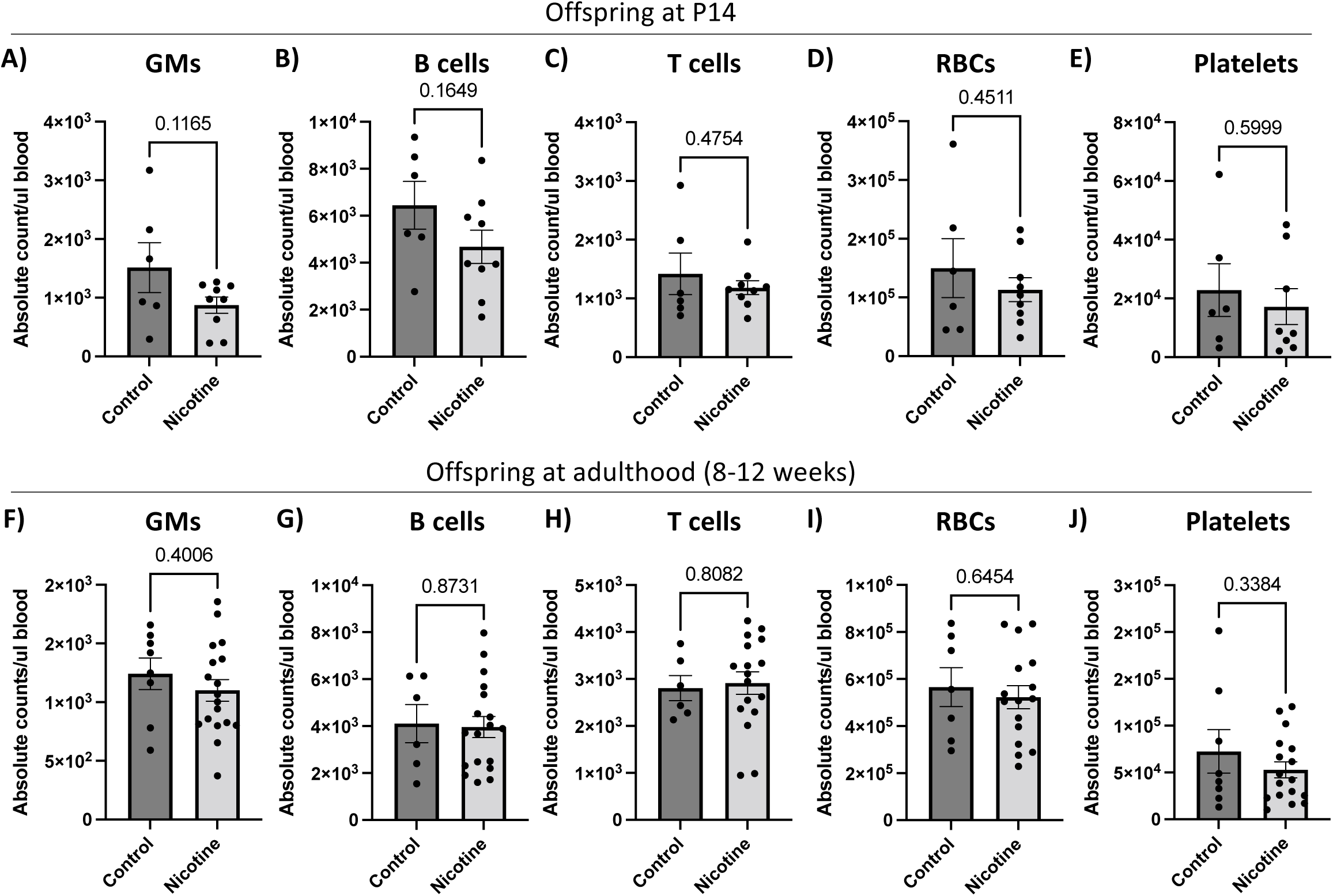
Perinatal nicotine exposure does not affect mature cell numbers in the peripheral blood. A-E) PNE does not alter mature cell numbers in the peripheral blood of P14 offspring. Quantification of absolute cell count per microliter of blood of “traditional” mature cells in the peripheral blood of P14 pups from control (n=6) and nicotine-exposed groups (n=9). Absolute cell counts of GMs (A), B cells (B), T cells (C), RBCs (D), and platelets (E). F-J) PNE does not alter mature cell numbers in the peripheral blood of adult offspring. Quantification of total numbers per microliter of “traditional” mature cells in the peripheral blood of adult offspring from control (n=8) and nicotine-exposed groups (n=18). Absolute cell counts of GMs (F), B cells (G), T cells (H), RBCs (I), and platelets (J). All errors bars represent SEM; each dot represents an individual animal.

### PNE leads to persistent decrease in tissue resident macrophages in the lungs

As smoking during pregnancy is strongly associated with higher susceptibility to respiratory diseases in the offspring, we wanted to determine if PNE results in persistent changes in establishment of non-traditional, tissue-resident immune cell populations that are established during the exposure period, and/or immune cells derived from PNE HSCs. Thus, we analyzed the lungs of pups born from nicotine-exposed mothers (**Figure 2A**) to be able to simultaneously compare traditional BM-derived and fetal-established immune cell populations (**Supplementary Figure 3A-C**). While there were no differences in lung weight being observed between PNE pups and controls (**Figure 4A**), the effects of PNE appeared to be cell-type specific in the lungs of adult mice. We observed no differences in cell numbers of CD4+ T cells (**Figure 4B**), CD8+ T cells (**Figure 4C**), TCRβ+ T cells (**Figure 4D**), TCRγδ T cells (**Figure 4E**), regulatory B cells (B regs; **Figure 4F**), neutrophils (**Figure 4G**) and eosinophils (**Figure 4H**). However, *in utero* nicotine exposure resulted in a long-lasting decrease of alveolar macrophages (**Figure 4I**) and interstitial monocytes/macrophages (**Figure 4J**). These data suggest that PNE primarily affects tissue-resident myeloid immune cell types that are established in the lung during fetal development and sustained throughout life with little contribution from adult hematopoiesis.

**Figure 4.**
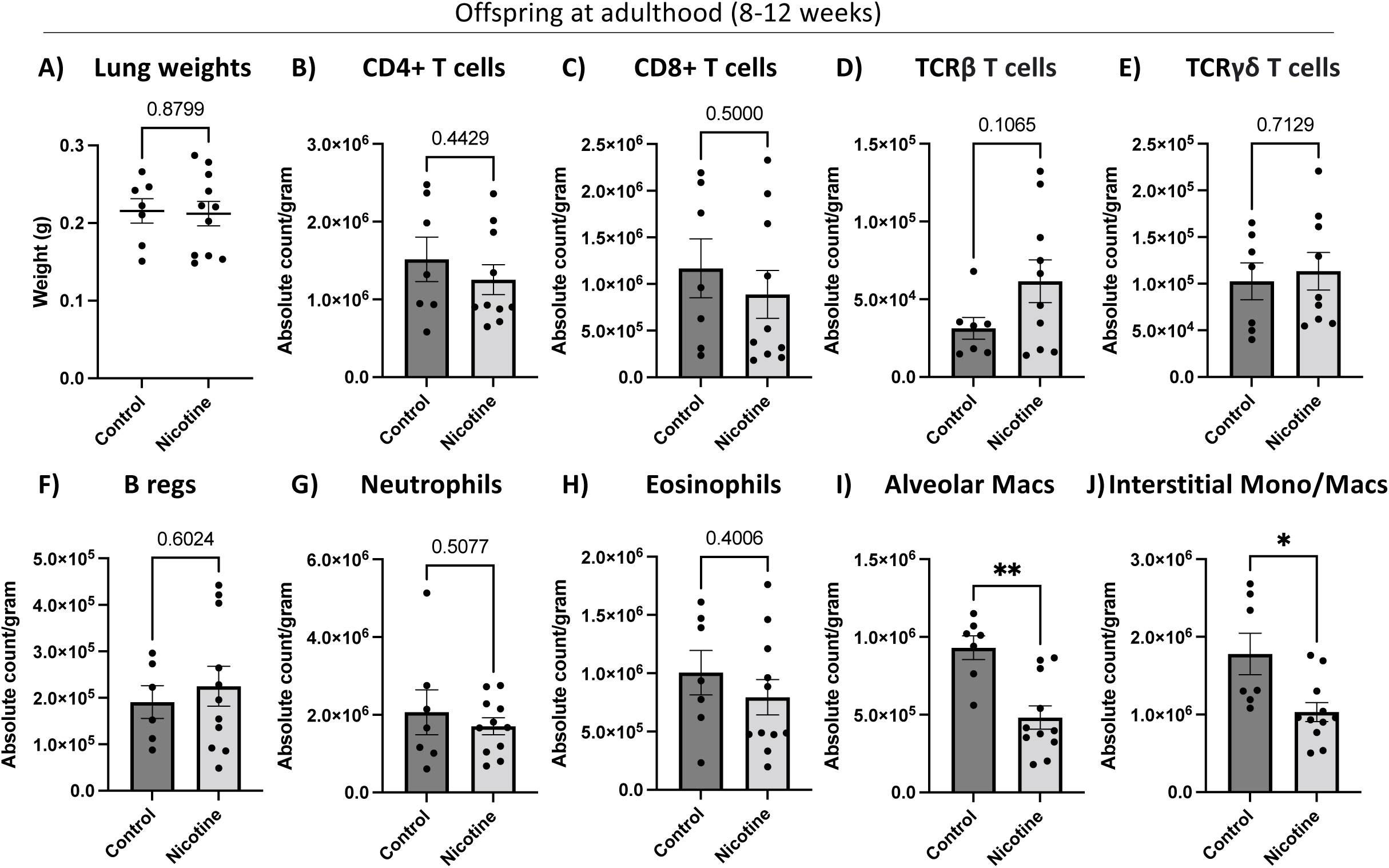
Perinatal nicotine exposure results in decreased macrophages in the adult lung. A) PNE does not alter the lung weight of adult offspring. Weight (g) of lungs from control (n=7) and nicotine-exposed (n=11) adult mice. B-J) PNE selectively affects alveolar and interstitial monocytes/macrophages of adult offsprings. Quantification of total numbers tissue resident immune cells in the lungs of the mice from Figure 4A. Absolute cell counts, normalized to grams of lung tissue, of CD3+ CD4+ T cells (B), CD3+ CD8+ T cells (C), TCRβ (D), TCRγδ T cells (E), B regs (F), neutrophils (G), eosinophils (H), alveolar macrophages (I), and interstitial monocytes/macrophages (J). *P<0.05, **P<0.01 (Student’s t-test). All errors bars represent SEM; each dot represents an individual animal.

### PNE exacerbates changes in immune cell proportions with secondary challenge

Children of smoking mothers are at significantly higher risk of infections. Having demonstrated that PNE permanently alters the number of phenotypic HSCs (**Figure 1-2**) and lung macrophages (**Figure 4**), we wanted to model this infection risk by determining how adult mice that had been perinatally exposed to nicotine would respond to a secondary immune challenge. To test this, we exposed the PNE adult offspring to a single high dose of lipopolysaccharide (LPS) and analyzed their BM and blood compartments 16 hours after injection (**Figure 5A**). The nucleated peripheral blood compartment of an adult mouse is composed of ∼75% lymphocytes (primarily B and T cells) and ∼25% myeloid cells (primarily GMs) (**Figure 5B**). LPS exposure alone resulted in decreased proportions of B and T cells, as well as an increase in the proportion of the GM pool in both control and PNE adult mice (**Figure 5B**). We observed a significantly greater increase in the proportion of GMs in mice that had been exposed to both nicotine perinatally and LPS in adulthood (**Figure 5B**). This was due to an overall increase in GM counts (**Figure 5C**), likely as a consequence of inflammation-induced accelerated granulopoiesis (Su et al., 2020), as well as a significant decrease in B and T cell numbers (**Figure 5D-E**), likely due to their movement out of the bloodstream and into tissues to fight off the perceived pathogen invasion. As expected, LPS-exposed animals showed significant expansion of the BM HSC pool and a significant depletion of MyPros in response to LPS (**Figure 5F-H**) (Zhang et al., 2016), while MPP numbers were unaffected (**Figure 5G**). The BM response to LPS appeared to be largely independent of previous nicotine exposure. In fact, there were no stark differences in the response to LPS between the control mice exposed to LPS only and PNE mice exposed to LPS for HSCs (**Figure 5F**), MPPs (**Figure 5G**), and MyPros (**Figure 5H**). Taken together, these data indicate that PNE may exacerbate the mature cell mediated response to a systemic immune trigger such as LPS; however, such potent inflammatory trigger might mask the extent to which PNE contributes to an altered response by BM HSPCs.

**Figure 5.**
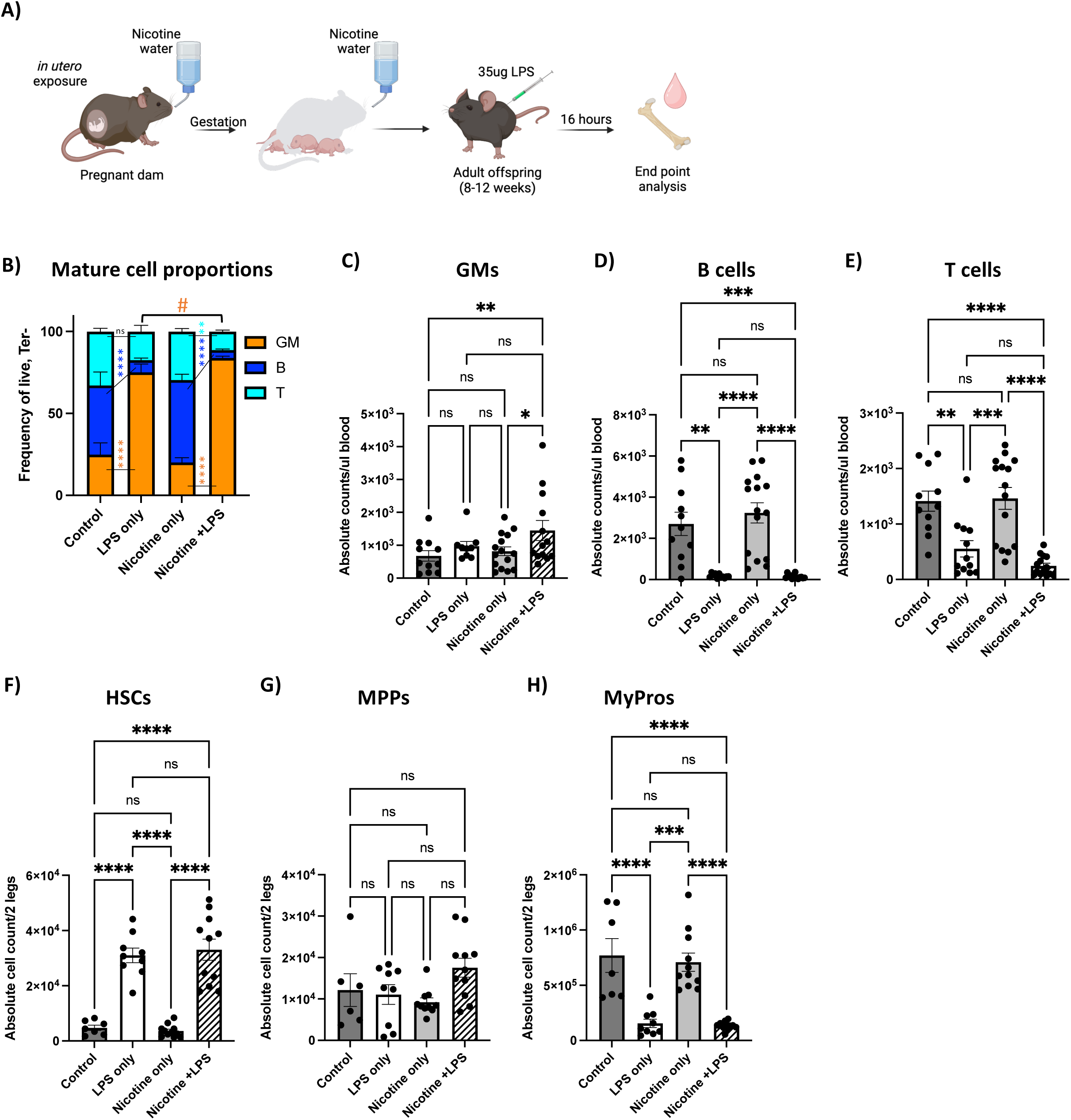
Perinatal nicotine exposure exacerbates emergency myelopoiesis in response to LPS. A) Experimental set up of LPS treatment. B) PNE significantly increases the proportion of peripheral blood GMs upon secondary LPS insult. Proportion of GM, B, and T cells in the peripheral blood of from control, LPS only, nicotine only, and nicotine +LPS. *P<0.05, **P<0.01, ****P<0.001 (Two-way ANOVA with Tukey post-hoc; only relevant comparisons shown). G-E) PNE exacerbates alterations in mature cell counts upon secondary LPS insult. Absolute cell counts of GMs (C), Bs (D), and Ts (E) from control (n=11), LPS only (n=12), nicotine only (n=15) and nicotine + LPS (n=14) mice. *P<0.05, **P<0.01, ***P<0.001, ****P<0.001 (Two-way ANOVA with Tukey post-hoc). F-H) PNE does not result in altered BM HSPC numbers upon secondary LPS insult compared to LPS only. Absolute cell counts of phenotypic HSCs (F), MPPs (G), and MyPros (H) from control (n=7), LPS only (n=9), nicotine only (n=11) and nicotine + LPS (n=11) mice. ***P<0.001, ****P<0.001 (Two-way ANOVA with Tukey post-hoc). Data represented as mean +/- SEM.

### Perinatal nicotine exposure leads to increased disease severity upon influenza infection

Given the previously reported link between maternal smoking and increased susceptibility to respiratory diseases in the offspring (Carroll et al., 2007; Cook & Strachan, 1999; DiFranza et al., 2004; Jaakkola et al., 2006; Jones et al., 2011; Taylor & Wadsworth, 1987), and our data showing a significant decrease in a lung macrophages in murine offspring of dams exposed to nicotine during gestation (**Figure 4**), we next investigated the effects of PNE to an acute viral respiratory infection. We infected adult PNE mice with mouse-adapted influenza A virus (IAV), commonly referred to as x31 [A/Aichi/02/68 (HA,NA) x A/Puerto Rico/8/34] (Sanders et al., 2013), which causes a non-lethal mild infection localized to the murine respiratory tract. To examine the response of PNE mice to IAV infection, we tracked their body weight and analyzed their blood counts during the IAV infection as well as BM and spleen 7 days post infection (dpi) (**Figure 6A**). Significant reductions in body weight started at 3 dpi IAV infection in all infected mice (**Figure 6B**). Interestingly, mice previously exposed to nicotine perinatally lost significantly more weight than PNE control infected mice starting at 4 dpi, indicative of increased severity of disease (**Figure 6B**). This was independent of sex and of body weight prior to infection (0 dpi) (**Supplementary Figure 4A**). Peripheral blood counts of both PNE and control mice exhibited a significant increase in CD11b+Gr1+ GMs at 3 dpi, which returned to baseline by 6 dpi (**Figure 6C**). Interestingly, PNE mice had significantly higher numbers of GMs, the majority of which are neutrophils, at 3 dpi compared to control mice infected with influenza (**Figure 6C**). Neutrophils are HSC-derived innate immune cells that were previously shown to increase in the lungs and blood after IAV infection and infiltrated into the respiratory tract (Camp & Jonsson, 2017) and that have been implicated in driving influenza disease severity (Brandes et al., 2013; Gautam et al., 2024). To confirm the identity of these peripheral blood neutrophils, we used an additional phenotypic marker combination, CD11b+ Ly6G+ Ly6C-, which led to similar outcomes (**Supplementary Figure 4G**). Regardless of PNE exposure, all infected mice developed lymphopenia (**Supplementary Figure 4C-D**) to similar levels. In contrast, PNE led to significantly more drastic reduction of natural killer (NK) cells in IAV-infected mice compared to non-exposed (control) IAV-infected mice (**Figure 6D**). The decrease in cytotoxic NK cells in the circulation may be explained by their sequestration to the lung, where they serve an important role in the immune response against influenza infection (Frank & Paust, 2020). RBC numbers remained unaffected upon IAV infection (**Supplementary Figure 4E**), while the number of platelets significantly increased at 6 dpi, independent of previous nicotine exposure, as recently also shown by others (Rommel et al., 2022) (**Supplementary Figure 4F**). PNE also caused exacerbated responses by immune cells in the lungs. Specifically, the alveolar macrophage pool, which was already substantially reduced in PNE mice (**Figure 4I**), was further diminished upon IAV infection in PNE mice compared to control mice infected with influenza (**Figure 6E**). The number of neutrophils in the lung, which was unaffected by PNE (**Figure 4G**), significantly increased upon IAV infection in PNE mice compared to control mice infected by influenza (**Figure 6F**). The significant increase in neutrophils and decrease in alveolar macrophages could be contributing to the increased disease severity indicated by the increased weight loss of PNE mice after IAV infection. Upon testing whether these changes originated in the BM, we did not observe significant changes in HSC numbers upon influenza infection (**Figure 6G**). In contrast, we found a significant decrease in MyPros, which include the hematopoietic progenitors of GMs, specifically in PNE mice infected with influenza compared to control mice infected with influenza (**Figure 6H**), potentially due to the significant demand for GMs (in particular neutrophils) during IAV infection. Overall, PNE led to increased disease susceptibility and severity upon IAV infection in adulthood.

**Figure 6.**
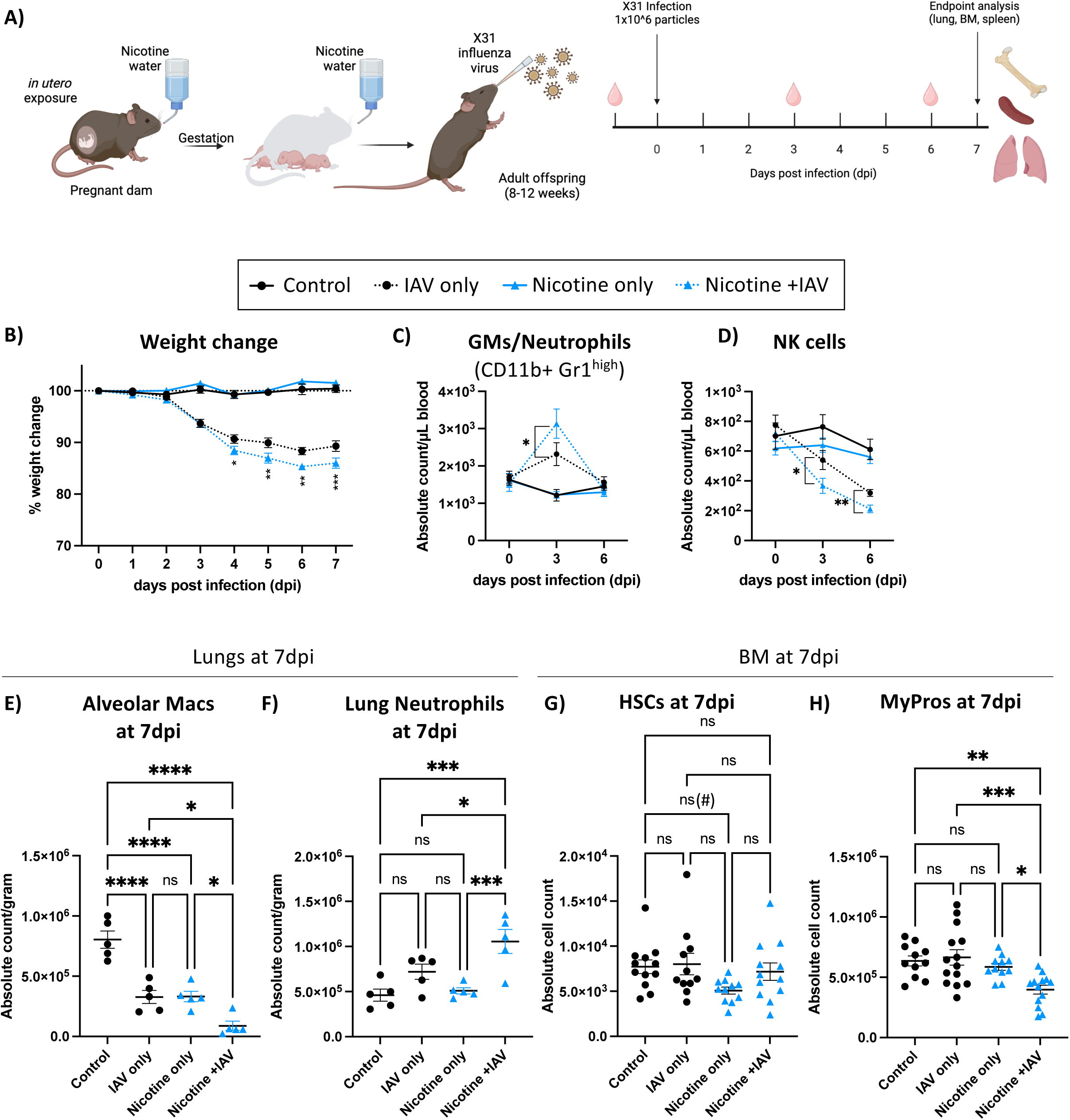
Perinatal nicotine exposure leads to increased disease severity upon influenza infection. A) Experimental set up of influenza (X31 IAV) treatment. B) PNE leads to significant weight loss upon IAV infection in adulthood. Weight loss as a percentage of original weight for the 4 treatment groups: Control + no IAV (n=16), Control + IAV (n=19), Nicotine + no IAV (n=16), and Nicotine + IAV (n=18). *P<0.05, **P<0.01, ***P<0.001 (Two-way ANOVA with Tukey’s post-hoc; only statistical differences between “Control + IAV” vs “Nicotine + IAV” are shown). C-D) PNE selectively alters peripheral blood immune cell numbers upon IAV infection. Quantification of absolute cell count per microliter of blood of GMs/neutrophils (C) and NK cells (D) at dpi 0, 3, and 6 (n=16-19/group). *P<0.05 (Two-way ANOVA with Tukey’s post-hoc; only statistical significance between infected groups shown). E-F) PNE significantly decreases alveolar macrophages and significantly increase lung neutrophils upon IAV infection. Absolute cell counts, normalized to grams of lung tissue, of alveolar macrophages (E) and neutrophils (F) at 7 dpi (n=5/group). *P<0.05, ***P<0.001, ****P<0.0001 (Two-way ANOVA with Tukey’s post-hoc). G-H) PNE results in significant loss of BM MyPros upon IAV. Absolute cell counts of BM HSCs (G) and MyPros (H) at 7 dpi (n=11-13/group). *P<0.05, **P<0.01, ***P<0.001 (Two-way ANOVA with Tukey’s post-hoc). (#) P<0.01 (Student’s t-test). All errors bars represent mean +/- SEM.

## DISCUSSION

The establishment of a robust immune system is crucial for lifelong health, enabling effective infection defense, response to challenges, and homeostasis. Disruptions during early immune development, such as exposure to toxicants like nicotine, can increase susceptibility to diseases, including autoimmune disorders and respiratory infections (Avşar et al., 2021; J. Cao et al., 2016; Cool et al., 2022; Dietz et al., 2010; Karthikeyan et al., 2024; Quelhas et al., 2018). Despite declining rates of maternal smoking during pregnancy, nicotine exposure remains a concern due to e-cigarettes and secondhand smoke. Here we showed that nicotine exposure during development perturbed immune establishment with long-term consequences. Mechanistically, this was manifested by altered cellular composition before and after subsequent infection. Trained immunity – characterized by epigenetic reprogramming of HSCs and innate immune cells – provides a framework to understand how perinatal exposures like nicotine may prime immune cells for altered responses later in life (R. Cao et al., 2024; de Laval et al., 2020; Divangahi et al., 2021; Johansson et al., 2023; Kain et al., 2023; Kaufmann et al., 2018; Netea et al., 2020).

Understanding the specific risks associated with PNE is necessary in addressing potential life-long health implications for the offspring. While it is well-documented that adult HSCs are impacted by inflammatory stimuli, including clinical influenza infections (Haas et al., 2015; Pietras et al., 2014; Rommel et al., 2022), our understanding of the effects of *in utero* exposure to toxic compounds on the establishment of the hematopoietic system remains limited. To shed light on this, our study focused on isolating the effects of nicotine alone, distinct from the diverse toxicants present in tobacco products. We observed a significant decrease in fetal and neonatal HSCs in PNE offspring (**Figure 1-2**), likely due to altered homing and migration patterns. This deficit persisted into adulthood (**Figure 2**). Notably, a reduction in HSC with PNE could not have been predicted by adult exposure, as nicotine treatment in adulthood led to the opposite result of increased HSC numbers (**Figure 2K-O**). Moreover, while adult nicotine caused elevated WBCs, the PNE-induced HSC reduction did not affect production of mature cells in the peripheral blood of adult PNE offspring (**Figure 3**) highlighting the remarkable capacity of hematopoietic progenitors to maintain hematopoietic homeostasis at steady-state.

Despite nearly normal steady-state hematopoiesis in adulthood, PNE exposure notably impacted the establishment of tissue-resident macrophages in the lungs (**Figure 4**) and primed the immune system for exaggerated responses. Nicotine-exposed mice showed increased disease severity during influenza infections (**Figure 6**), with greater weight loss, elevated neutrophil counts, and reduced alveolar macrophages. These results demonstrate that very selective cellular mechanisms provide resilience to infectious disease, and that, surprisingly, traditional adaptive immune cells were not evident key players. This observation may point to key mechanistic differences of fetal and adult exposures that are anchored in the dynamic regulation of immune system development. Our findings suggest that nicotine exposure may induce trained immunity by priming both HSCs and innate immune cells for maladaptive responses to respiratory infections later in life. This mechanism aligns with established links between maternal smoking and increased respiratory infection susceptibility in humans (Carroll et al., 2007; Cook & Strachan, 1999; DiFranza et al., 2004; Jaakkola et al., 2006; Jones et al., 2011; Taylor & Wadsworth, 1987). Further studies are needed to elucidate how nicotine-induced epigenetic and metabolic changes influence trained immunity and immune dysfunction. Such insights have the potential to inform therapeutic strategies to mitigate long-term health impacts of maternal exposure to nicotine and other toxins.

## ACKNOWLEDGMENTS

This work was supported by Tobacco Disease Research Program (TRDRP) predoctoral fellowships to T.C. and A.R.y.B; UCSC Dissertation Year Fellowship to A.R.y.B; NIH National Institute of General Medical Sciences (NIGMS) IRACDA (K12GM139185) to A.R.y.B. and B.A.M. NICHD T32 (T32HD108079) to B.A.M. and matching funds to S.C., CIRM predoctoral fellowship to S.S. (EDUC4-12759) and postdoctoral fellowship to M.G.E.R. CIRM Bridges S.C.I.L.L. to A.D. NIDDK and NIA awards (R01DK100917 and R01AG062879) to E.C.F, CIRM Facilities awards CL1-00506 and FA1-00617-1 (RRIDs SCR_021149, SCR_021353 and SCR_021135) to University of California, Santa Cruz. Some images were generated using BioRender license number #2617-2694.

## METHODS

### Mice

All animals were housed and bred in the AALAC accredited vivarium at UC Santa Cruz and group housed in ventilated cages on a standard 12:12 light cycle. All procedures were approved by the UCSC or the UC Merced Institutional Animal Care and Use (IACUC) committees. WT C57BL/6 mice were used for all experiments. Male and female mice were used equally and without sex discrimination for all experiments. Mice were fed normal chow diet and given nicotine solution (100 µg/mL nicotine and 5% sucrose, diluted in water(Maier et al., 2011)) *ad libitum*. Mice were sacrificed at post-natal day 0 (P0), post-natal day 14 (P14), or adulthood (8-12 weeks of age). For LPS exposure, adult mice (8-12 weeks) were administered a single intraperitoneal injection of 35 µg of LPS. They were sacrificed for analysis ∼16 hours post injection.

### Tissue and cell isolation

Mice were sacrificed by CO_2_ inhalation. Neonatal livers were harvested and homogenized using mortar and pestle, or microscope slides, before filtering through a 70 µm filter. Lungs were harvested and treated with 1X PBS (Ca+/Mg+) with 2% fetal bovine serum (WVR) and 2 mg/mL Collagenase IV (Gibco) and 100 U/mL DNaseI (Sigma) for 1 hour at 37°C. Following incubation, tissue was passed through a 16G needle followed by 19G needle (approximately 10 times each) and then filtered through a 70 µm filter. For all BM HSPC analysis, both long bones (tibia and femur) were pulled, crushed in 1X PBS supplemented with 5 mM EDTA with 2% fetal bovine serum, and single cell suspensions were filtered through a 70 µm filter. Peripheral blood (PB) taken from the femoral artery at sacrifice for terminal analysis or by retro-orbital bleeds for infection experiments.

### Cytokine analysis

Livers were extracted from P0 pups, weighed, and homogenized in 200µL of PBS without calcium and magnesium to collect serum. After a 10-minute centrifugation at 300xg at room temperature, 180 µL of supernatant were transferred to a new tube, incubated at room temperature for 30 minutes (to allow clotting), centrifuged at 7000xg for 7min at 4C, and finally 75 µL of the supernatant was sent to Eve’s Technologies for “Mouse Cytokine/Chemokine 44-Plex Discovery Assay Array”. Concentrations of these cytokines/chemokines in the liver serum was normalized to total protein concentrations determined by Eve’s Technologies.

### Flow cytometry

Single cell suspensions were stained with monoclonal anti-mouse antibodies on ice in the dark for 20 minutes and acquired using a FACSAria, LSRII flow cytometer (BD Biosciences, San Jose, CA) or Cytoflex LX (Beckman Coulter) at the University of California-Santa Cruz, as described previously (Beaudin et al., 2014, 2016; Boyer et al., 2011; Cool et al., 2020; Poscablo et al., 2021; Rajendiran et al., 2020; Rodriguez Y Baena et al., 2022; Smith-Berdan et al., 2015; Worthington et al., 2022). Cell populations were defined by the following cell surface markers: P0 liver HSC (Live, CD3-CD4-CD5-CD8-B220-Gr1-Ter119-cKit+Sca1+CD150+); P14 and adult BM HSCs (Live, CD3-CD4-CD5-CD8-B220-Gr1-CD11b-Ter119-cKit+Sca1+CD150+ Flk2-); BM MPP (Live, CD3-CD4-CD5-CD8-B220-Gr1-CD11b-Ter119-cKit+Sca1+Flk2+CD150-); BM MyPro (Live, CD3-CD4-CD5-CD8-B220-Gr1-CD11b-Ter119-cKit+Sca1-); BM CMP (Live, CD3-CD4-CD5- CD8-B220-Gr1-CD11b-Ter119-cKit+Sca1-CD16/32midCD34+); BM GMP (Live, CD3-CD4-CD5- CD8-B220-Gr1-CD11b-Ter119-cKit+Sca1-CD16/32+CD34+); BM MEP (Live, CD3-CD4-CD5- CD8-B220-Gr1-CD11b-Ter119-cKit+Sca1-CD16/32-CD34-); PB GM (Live, Ter119-CD61-B220- CD3-CD11b+Gr1+), PB neutrophils (Live, Ter119-CD61-B220-CD3-CD11b+ Gr1^high^ or Live, Ter119-CD61-B220-CD3-CD11b+Ly6G+Ly6C-), PB B cells (Live, Ter119-CD61-CD11b-Gr1- B220+CD3-), PB T cells (Live, Ter119-CD61-CD11b-Gr1-B220-CD3+), PB RBC (Live, CD61- B220-CD3-Ter119+), PB platelets (Live, CD61+B220-CD3-Ter119-), PB NK cells (Live, Ter119- CD45+B220-Gr1-NK1.1+), lung CD4+ T cells (Live, Ter119-Gr1-B220-CD11b-CD3+CD8-CD4+), lung CD8+ T cells (Live, Ter119-Gr1-B220-CD11b-CD3+CD4-CD8+), lung TCRβ T cells (Live, Ter119-Gr1-B220-CD11b-CD3+CD4-CD8-TCRγδ-TCRβ+), lung TCRγδ T cells (Live, Ter119- Gr1-B220-CD11b-CD3+CD4-CD8-TCRβ-TCRγδ+), lung B regs (Live, CD3-CD4-CD8-Ter119- Gr1-CD11b-CD19+IgM+CD21+CD23+), lung alveolar macrophages (Live, CD45+CD11b- CD11c+Ly6G-SiglecF+), lung interstitial macrophages (Live, CD45+CD11b+CD11c+Ly6G- SiglecF-), lung eosinophils (Live, CD3-CD4-CD5-CD8-B220-Ter119-CD45+CD11b+CD11c- Ly6G-SiglecF+), lung neutrophils (Live, CD3-CD4-CD5-CD8-B220-Ter119- CD45+CD11b+CD11c-Ly6G+SiglecF). Data was analyzed using FlowJo 10.10.

### Cell cycle analysis

Single cell suspension from P0 neonatal liver and P14 BM were first stained with lineage markers (Ter119, Gr1, B220, CD3, CD4, CD5, CD8) and HSPC markers (cKit, Sca1, CD150, Flk2), then fixed with 4% PFA, permeabilized with 0.3% saponin, treated with 100U RNAase A (ThermoFisher), and stained with 0.01 mg/ml DAPI (Millipore) or Hoechst 33342 (ThermoFisher) for cell cycle analysis by flow cytometry.

### qPCR of nAChRa7

For whole tissue samples, total RNA was isolated, crushed brain and liver with a Direct-zol RNA MiniPrep kit (Zymo Research). For sorted cells, RNA was isolated from twenty thousand purified fetal liver HSCs using Trizol (Life Technologies) and a DNase treatment step. Complementary DNA (cDNA) was synthesized using the High-Capacity cDNA Reverse Transcription Kit (ThermoFisher). Quantitative PCR was performed using a Viia 7 Real-Time PCR (Applied Biosystems). Fold expression relative to the reference gene (GAPDH) was calculated using the comparative CT method (ΔΔCT) and the values were normalized to the positive control (brain tissue). The primers used included: GAPDH - Forward: 5’-TGTGTCCGTCGTGGATCTGA-3’; GAPDH - Reverse: 5’-CCTGCTTCACCACCTTCTTGA-3’; nAChRa7 - Forward: 5’- TTGTGCTGCGATATCACCAC-3’; nAChRa7 - Reverse: 5’TTCATGCGCAGAAACCATGC-3’.

### Influenza Infections

Prior to influenza infection, mice were anesthetized by isoflurane inhalation, and then inoculated intranasally (Sanders et al., 2013) in both nostrils with 10^6 egg infection dose 50 (EID_50_) of influenza H3N2 virus A/Aichi/02/68 (HA, NA) x A/Puerto Rico/8/34 (diluted in DPBS) in 30 µL or 30 µL of 1x DPBS using a displacement pipet. Mice were monitored post-inoculation to ensure recovery from anesthesia. As per our institutionally approved IACUC protocol, mice underwent daily monitoring of body weight and were euthanized following evidence (body index score) of severe morbidity.

### Quantification and statistical analysis

Unpaired two-tailed Student’s t-tests and ANOVAs adjusted for multiple comparisons with Tukey or Dunnett’s post-hoc tests were used to assess statistical significance for comparisons of different groups, as appropriate. The sample size (n) and p values are provided for each experiment in the respective figure legend. For each figure, at least 2-3 independent experiments were performed. All data are shown as mean ±. S.E.M unless stated otherwise. Outlier analysis tests were performed, and data points were removed subsequently as appropriate.

## REFERENCES

Apostol, A. C., Jensen, K. D. C., & Beaudin, A. E. (2020). Training the Fetal Immune System Through Maternal Inflammation—A Layered Hygiene Hypothesis. Frontiers in Immunology, 11, 123. 10.3389/fimmu.2020.00123

Avşar, T. S., McLeod, H., & Jackson, L. (2021). Health outcomes of smoking during pregnancy and the postpartum period: An umbrella review. BMC Pregnancy and Childbirth, 21(1), 254. 10.1186/s12884-021-03729-1

Beaudin, A. E., Boyer, S. W., & Forsberg, E. C. (2014). Flk2/Flt3 promotes both myeloid and lymphoid development by expanding non-self-renewing multipotent hematopoietic progenitor cells. Experimental Hematology, 42(3), 218–229.e4. 10.1016/j.exphem.2013.11.013

Beaudin, A. E., Boyer, S. W., Perez-Cunningham, J., Hernandez, G. E., Derderian, S. C., Jujjavarapu, C., Aaserude, E., MacKenzie, T., & Forsberg, E. C. (2016). A transient developmental hematopoietic stem cell gives rise to innate-like B and T cells. Cell Stem Cell, 19(6), 768–783. 10.1016/j.stem.2016.08.013

Ben-Yehudah, A., Campanaro, B. M., Wakefield, L. M., Kinney, T. N., Brekosky, J., Eisinger, V. M., Castro, C. A., & Carlisle, D. L. (2013). Nicotine exposure during differentiation causes inhibition of N-myc expression. Respiratory Research, 14(1), 119. 10.1186/1465-9921-14-119

Boisset, J.-C., Van Cappellen, W., Andrieu-Soler, C., Galjart, N., Dzierzak, E., & Robin, C. (2010). In vivo imaging of haematopoietic cells emerging from the mouse aortic endothelium. Nature, 464(7285), 116–120. 10.1038/nature08764

Boyer, S. W., Schroeder, A. V., Smith-Berdan, S., & Forsberg, E. C. (2011). All hematopoietic cells develop from hematopoietic stem cells through Flk2/Flt3-positive progenitor cells. Cell Stem Cell, 9(1), 64. 10.1016/j.stem.2011.04.021

Brandes, M., Klauschen, F., Kuchen, S., & Germain, R. N. (2013). A systems analysis identifies a feedforward inflammatory circuit leading to lethal influenza infection. Cell, 154(1), 197–212. 10.1016/j.cell.2013.06.013

Camp, J. V., & Jonsson, C. B. (2017). A Role for Neutrophils in Viral Respiratory Disease. Frontiers in Immunology, 8, 550. 10.3389/fimmu.2017.00550

Cao, J., Xu, X., Hylkema, M. N., Zeng, E. Y., Sly, P. D., Suk, W. A., Bergman, Å., & Huo, X. (2016). Early-life Exposure to Widespread Environmental Toxicants and Health Risk: A Focus on the Immune and Respiratory Systems. Annals of Global Health, 82(1), 119. 10.1016/j.aogh.2016.01.023

Cao, R., Thatavarty, A., & King, K. Y. (2024). Forged in the fire: Lasting impacts of inflammation on hematopoietic progenitors. Experimental Hematology, 134, 104215. 10.1016/j.exphem.2024.104215

Carroll, K. N., Gebretsadik, T., Griffin, M. R., Dupont, W. D., Mitchel, E. F., Wu, P., Enriquez, R., & Hartert, T. V. (2007). Maternal Asthma and Maternal Smoking Are Associated With Increased Risk of Bronchiolitis During Infancy. Pediatrics, 119(6), 1104–1112. 10.1542/peds.2006-2837

Chang, E., Forsberg, E. C., Wu, J., Bingyin Wang, Prohaska, S. S., Allsopp, R., Weissman, I. L., & Cooke, J. P. (2010). Cholinergic activation of hematopoietic stem cells: Role in tobacco-related disease? Vascular Medicine, 15(5), 375–385. 10.1177/1358863X10378377

Collins, A. C., Pogun, S., Nesil, T., & Kanit, L. (2012). Oral Nicotine Self-Administration in Rodents. *Journal of Addiction Research & Therapy*, S2, 004. 10.4172/2155-6105.S2-004

Collins, A., Swann, J. W., Proven, M. A., Patel, C. M., Mitchell, C. A., Kasbekar, M., Dellorusso, P. V., & Passegué, E. (2024). Maternal inflammation regulates fetal emergency myelopoiesis. Cell, 187(6), 1402–1421.e21. 10.1016/j.cell.2024.02.002

Cook, D. G., & Strachan, D. P. (1999). Health effects of passive smoking 10: Summary of effects of parental smoking on the respiratory health of children and implications for research. Thorax, 54(4), 357–366. 10.1136/thx.54.4.357

Cool, T., Baena, A. R., & Forsberg, E. C. (2022). Clearing the Haze: How Does Nicotine Affect Hematopoiesis before and after Birth? Cancers, 14(1). 10.3390/cancers14010184

Cool, T., & Forsberg, E. C. (2019). Chasing Mavericks: The quest for defining developmental waves of hematopoiesis. Current Topics in Developmental Biology, 132, 1–29. 10.1016/bs.ctdb.2019.01.001

Cool, T., Worthington, A., Poscablo, D., Hussaini, A., & Forsberg, E. C. (2020). Interleukin 7 receptor is required for myeloid cell homeostasis and reconstitution by hematopoietic stem cells. Experimental Hematology, 90, 39–45.e3. 10.1016/j.exphem.2020.09.001

de Laval, B., Maurizio, J., Kandalla, P. K., Brisou, G., Simonnet, L., Huber, C., Gimenez, G., Matcovitch-Natan, O., Reinhardt, S., David, E., Mildner, A., Leutz, A., Nadel, B., Bordi, C., Amit, I., Sarrazin, S., & Sieweke, M. H. (2020). C/EBPβ-Dependent Epigenetic Memory Induces Trained Immunity in Hematopoietic Stem Cells. Cell Stem Cell, 26(5), 657–674.e8. 10.1016/j.stem.2020.01.017

Dietz, P. M., England, L. J., Shapiro-Mendoza, C. K., Tong, V. T., Farr, S. L., & Callaghan, W. M. (2010). Infant morbidity and mortality attributable to prenatal smoking in the U.S. American Journal of Preventive Medicine, 39(1), 45–52. 10.1016/j.amepre.2010.03.009

DiFranza, J. R., Aligne, C. A., & Weitzman, M. (2004). Prenatal and postnatal environmental tobacco smoke exposure and children’s health. Pediatrics, *113*(4 Suppl), 1007–1015.

Divangahi, M., Aaby, P., Khader, S. A., Barreiro, L. B., Bekkering, S., Chavakis, T., van Crevel, R., Curtis, N., DiNardo, A. R., Dominguez-Andres, J., Duivenvoorden, R., Fanucchi, S., Fayad, Z., Fuchs, E., Hamon, M., Jeffrey, K. L., Khan, N., Joosten, L. A. B., Kaufmann, E., … Netea, M. G. (2021). Trained immunity, tolerance, priming and differentiation: Distinct immunological processes. Nature Immunology, 22(1), 2–6. 10.1038/s41590-020-00845-6

Frank, K., & Paust, S. (2020). Dynamic Natural Killer Cell and T Cell Responses to Influenza Infection. Frontiers in Cellular and Infection Microbiology, 10, 425. 10.3389/fcimb.2020.00425

Gautam, A., Boyd, D. F., Nikhar, S., Zhang, T., Siokas, I., Van de Velde, L.-A., Gaevert, J., Meliopoulos, V., Thapa, B., Rodriguez, D. A., Cai, K. Q., Yin, C., Schnepf, D., Beer, J., DeAntoneo, C., Williams, R. M., Shubina, M., Livingston, B., Zhang, D., … Balachandran, S. (2024). Necroptosis blockade prevents lung injury in severe influenza. Nature, 628(8009), 835–843. 10.1038/s41586-024-07265-8

Gracia, M. C. (2005). Exposure to nicotine is probably a major cause of inflammatory diseases among non-smokers. Medical Hypotheses, 65(2), 253–258. 10.1016/j.mehy.2005.02.039

Guilliams, M., De Kleer, I., Henri, S., Post, S., Vanhoutte, L., De Prijck, S., Deswarte, K., Malissen, B., Hammad, H., & Lambrecht, B. N. (2013). Alveolar macrophages develop from fetal monocytes that differentiate into long-lived cells in the first week of life via GM-CSF. Journal of Experimental Medicine, 210(10), 1977–1992. 10.1084/jem.20131199

Haas, S., Hansson, J., Klimmeck, D., Loeffler, D., Velten, L., Uckelmann, H., Wurzer, S., Prendergast, Á. M., Schnell, A., Hexel, K., Santarella-Mellwig, R., Blaszkiewicz, S., Kuck, A., Geiger, H., Milsom, M. D., Steinmetz, L. M., Schroeder, T., Trumpp, A., Krijgsveld, J., & Essers, M. A. G. (2015). Inflammation-Induced Emergency Megakaryopoiesis Driven by Hematopoietic Stem Cell-like Megakaryocyte Progenitors. Cell Stem Cell, 17(4), 422–434. 10.1016/j.stem.2015.07.007

Hildreth, A. D., & O’Sullivan, T. E. (2019). Tissue-Resident Innate and Innate-Like Lymphocyte Responses to Viral Infection. Viruses, 11(3), 272. 10.3390/v11030272

Isotani, R., Igarashi, M., Miura, M., Naruse, K., Kuranami, S., Katoh, M., Nomura, S., & Yamauchi, T. (2025). Nicotine enhances the stemness and tumorigenicity in intestinal stem cells via Hippo-YAP/TAZ and Notch signal pathway. eLife, 13, RP95267. 10.7554/eLife.95267.4

Jaakkola, J. J., Kosheleva, A. A., Katsnelson, B. A., Kuzmin, S. V., Privalova, L. I., & Spengler, J. D. (2006). Prenatal and postnatal tobacco smoke exposure and respiratory health in Russian children. Respiratory Research, 7(1), 48. 10.1186/1465-9921-7-48

Johansson, A., Lin, D. S., Mercier, F. E., Yamashita, M., Divangahi, M., & Sieweke, M. H. (2023). Trained immunity and epigenetic memory in long-term self-renewing hematopoietic cells. Experimental Hematology, 121, 6–11. 10.1016/j.exphem.2023.02.001

Jones, L. L., Hashim, A., McKeever, T., Cook, D. G., Britton, J., & Leonardi-Bee, J. (2011). Parental and household smoking and the increased risk of bronchitis, bronchiolitis and other lower respiratory infections in infancy: Systematic review and meta-analysis. Respiratory Research, 12(1), 5. 10.1186/1465-9921-12-5

Kain, B. N., Tran, B. T., Luna, P. N., Cao, R., Le, D. T., Florez, M. A., Maneix, L., Toups, J. D., Morales-Mantilla, D. E., Koh, S., Han, H., Jaksik, R., Huang, Y., Catic, A., Shaw, C. A., & King, K. Y. (2023). Hematopoietic stem and progenitor cells confer cross-protective trained immunity in mouse models. iScience, 26(9), 107596. 10.1016/j.isci.2023.107596

Karthikeyan, B. S., Hyötyläinen, T., Ghaffarzadegan, T., Triplett, E., Orešič, M., & Ludvigsson, J. (2024). Prenatal exposure to environmental contaminants and cord serum metabolite profiles in future immune-mediated diseases. Journal of Exposure Science & Environmental Epidemiology, 34(4), 647–658. 10.1038/s41370-024-00680-z

Katono, T., Kawato, T., Tanabe, N., Suzuki, N., Yamanaka, K., Oka, H., Motohashi, M., & Maeno, M. (2006). Nicotine Treatment Induces Expression of Matrix Metalloproteinases in Human Osteoblastic Saos-2 Cells. Acta Biochimica et Biophysica Sinica, 38(12), 874–882. 10.1111/j.1745-7270.2006.00240.x

Kaufmann, E., Sanz, J., Dunn, J. L., Khan, N., Mendonça, L. E., Pacis, A., Tzelepis, F., Pernet, E., Dumaine, A., Grenier, J.-C., Mailhot-Léonard, F., Ahmed, E., Belle, J., Besla, R., Mazer, B., King, I. L., Nijnik, A., Robbins, C. S., Barreiro, L. B., & Divangahi, M. (2018). BCG Educates Hematopoietic Stem Cells to Generate Protective Innate Immunity against Tuberculosis. Cell, 172(1–2), 176–190.e19. 10.1016/j.cell.2017.12.031

Kizildag, S., Hosgorler, F., Güvendi, G., Koc, T. B., Kandis, S., Argon, A., Ates, M., & Uysal, N. (2021). Nicotine lowers TNF-α, IL-1b secretion and leukocyte accumulation via nAChR in rat stomach. Toxin Reviews, 40(1), 17–24. 10.1080/15569543.2020.1790604

Konno, K., Sasaki, T., Kulkeaw, K., & Sugiyama, D. (2020). Paracrine CCL17 and CCL22 signaling regulates hematopoietic stem/progenitor cell migration and retention in mouse fetal liver. Biochemical and Biophysical Research Communications, 527(3), 730–736. 10.1016/j.bbrc.2020.04.045

Lazarov, T., Juarez-Carreño, S., Cox, N., & Geissmann, F. (2023). Physiology and diseases of tissue-resident macrophages. Nature, 618(7966), 698–707. 10.1038/s41586-023-06002-x

Lewis, K., Yoshimoto, M., & Takebe, T. (2021). Fetal liver hematopoiesis: From development to delivery. Stem Cell Research & Therapy, 12(1), 139. 10.1186/s13287-021-02189-w

Li, Q., Zhou, X.-D., Kolosov, V. P., & Perelman, J. M. (2011). Nicotine reduces TNF-α expression through a α7 nAChR/MyD88/NF-ĸB pathway in HBE16 airway epithelial cells. *Cellular Physiology and Biochemistry: International Journal of Experimental Cellular Physiology*, Biochemistry, and Pharmacology, 27(5), 605–612. 10.1159/000329982

Lichanska, A. M., & Hume, D. A. (2000). Origins and functions of phagocytes in the embryo. Experimental Hematology, 28(6), 601–611. 10.1016/s0301-472x(00)00157-0

López, D. A., Otsuka, K. S., Apostol, A. C., Posada, J., Sánchez-Arcila, J. C., Jensen, K. D., & Beaudin, A. E. (2023). Both maternal IFNγ exposure and acute prenatal infection with Toxoplasma gondii activate fetal hematopoietic stem cells. The EMBO Journal, 42(14), e112693. 10.15252/embj.2022112693

Maier, C. R., Hollander, M. C., Hobbs, E. A., Dogan, I., Linnoila, R. I., & Dennis, P. A. (2011). Nicotine Does Not Enhance Tumorigenesis in Mutant *K*—*Ras* –Driven Mouse Models of Lung Cancer. Cancer Prevention Research, 4(11), 1743–1751. 10.1158/1940-6207.CAPR-11-0365

Mohsenzadeh, Y., Rahmani, A., Cheraghi, J., Pyrani, M., & Asadollahi, K. (2014). Prenatal exposure to nicotine in pregnant rat increased inflammatory marker in Newborn Rat. Mediators of Inflammation, 2014. 10.1155/2014/274048

Netea, M. G., Domínguez-Andrés, J., Barreiro, L. B., Chavakis, T., Divangahi, M., Fuchs, E., Joosten, L. A. B., Van Der Meer, J. W. M., Mhlanga, M. M., Mulder, W. J. M., Riksen, N. P., Schlitzer, A., Schultze, J. L., Stabell Benn, C., Sun, J. C., Xavier, R. J., & Latz, E. (2020). Defining trained immunity and its role in health and disease. Nature Reviews Immunology, 20(6), 375–388. 10.1038/s41577-020-0285-6

Pattenden, S., Antova, T., Neuberger, M., Nikiforov, B., De Sario, M., Grize, L., Heinrich, J., Hruba, F., Janssen, N., Luttmann-Gibson, H., Privalova, L., Rudnai, P., Splichalova, A., Zlotkowska, R., & Fletcher, T. (2006). Parental smoking and children’s respiratory health: Independent effects of prenatal and postnatal exposure. Tobacco Control, 15(4), 294–301. 10.1136/tc.2005.015065

Piao, W.-H., Campagnolo, D., Dayao, C., Lukas, R. J., Wu, J., & Shi, F.-D. (2009). Nicotine and inflammatory neurological disorders. Acta Pharmacologica Sinica, 30(6), 715–722. 10.1038/aps.2009.67

Pietras, E. M., Lakshminarasimhan, R., Techner, J.-M., Fong, S., Flach, J., Binnewies, M., & Passegué, E. (2014). Re-entry into quiescence protects hematopoietic stem cells from the killing effect of chronic exposure to type I interferons. Journal of Experimental Medicine, 211(2), 245–262. 10.1084/jem.20131043

Poscablo, D. M., Worthington, A. K., Smith-Berdan, S., & Forsberg, E. C. (2021). Megakaryocyte progenitor cell function is enhanced upon aging despite the functional decline of aged hematopoietic stem cells. Stem Cell Reports, 16(6), 1598–1613. 10.1016/j.stemcr.2021.04.016

Quelhas, D., Kompala, C., Wittenbrink, B., Han, Z., Parker, M., Shapiro, M., Downs, S., Kraemer, K., Fanzo, J., Morris, S., & Kreis, K. (2018). The association between active tobacco use during pregnancy and growth outcomes of children under five years of age: A systematic review and meta-analysis. BMC Public Health, 18(1), 1372. 10.1186/s12889-018-6137-7

Rajendiran, S., Boyer, S. W., & Forsberg, E. C. (2020). A quantitative hematopoietic stem cell reconstitution protocol: Accounting for recipient variability, tissue distribution and cell half-lives. Stem Cell Research, 50, 102145. 10.1016/j.scr.2020.102145

Rodriguez Y Baena, A., Rajendiran, S., Manso, B. A., Krietsch, J., Boyer, S. W., Kirschmann, J., & Forsberg, E. C. (2022). New transgenic mouse models enabling pan-hematopoietic or selective hematopoietic stem cell depletion in vivo. Scientific Reports, 12(1), 3156. 10.1038/s41598-022-07041-6

Rommel, M. G. E., Walz, L., Fotopoulou, F., Kohlscheen, S., Schenk, F., Miskey, C., Botezatu, L., Krebs, Y., Voelker, I. M., Wittwer, K., Holland-Letz, T., Ivics, Z., Von Messling, V., Essers, M. A. G., Milsom, M. D., Pfaller, C. K., & Modlich, U. (2022). Influenza A virus infection instructs hematopoiesis to megakaryocyte-lineage output. Cell Reports, 41(1), 111447. 10.1016/j.celrep.2022.111447

Sanders, C. J., Vogel, P., McClaren, J. L., Bajracharya, R., Doherty, P. C., & Thomas, P. G. (2013). Compromised respiratory function in lethal influenza infection is characterized by the depletion of type I alveolar epithelial cells beyond threshold levels. American Journal of Physiology-Lung Cellular and Molecular Physiology, 304(7), L481–L488. 10.1152/ajplung.00343.2012

Schaal, C., & Chellappan, S. P. (2014). Nicotine-Mediated Cell Proliferation and Tumor Progression in Smoking-Related Cancers. Molecular Cancer Research, 12(1), 14–23. 10.1158/1541-7786.MCR-13-0541

Smith-Berdan, S., Nguyen, A., Hong, M. A., & Forsberg, E. C. (2015). ROBO4-mediated vascular integrity regulates the directionality of hematopoietic stem cell trafficking. Stem Cell Reports, 4(2), 255–268. 10.1016/j.stemcr.2014.12.013

Sorokin, S. P., Hoyt, R. F., Blunt, D. G., & McNelly, N. A. (1992). Macrophage development: II. Early ontogeny of macrophage populations in brain, liver, and lungs of rat embryos as revealed by a lectin marker. The Anatomical Record, 232(4), 527–550. 10.1002/ar.1092320410

Su, Y., Gao, J., Kaur, P., & Wang, Z. (2020). Neutrophils and Macrophages as Targets for Development of Nanotherapeutics in Inflammatory Diseases. Pharmaceutics, 12(12), 1222. 10.3390/pharmaceutics12121222

Taylor, B., & Wadsworth, J. (1987). Maternal smoking during pregnancy and lower respiratory tract illness in early life. Archives of Disease in Childhood, 62(8), 786–791. 10.1136/adc.62.8.786

Valdez-Miramontes, C. E., Trejo Martínez, L. A., Torres-Juárez, F., Rodríguez Carlos, A., Marin-Luévano, S. P., de Haro-Acosta, J. P., Enciso-Moreno, J. A., & Rivas-Santiago, B. (2020). Nicotine modulates molecules of the innate immune response in epithelial cells and macrophages during infection with M. tuberculosis. Clinical and Experimental Immunology, 199(2), 230–243. 10.1111/cei.13388

Vivier, E., Artis, D., Colonna, M., Diefenbach, A., Di Santo, J. P., Eberl, G., Koyasu, S., Locksley, R. M., McKenzie, A. N. J., Mebius, R. E., Powrie, F., & Spits, H. (2018). Innate Lymphoid Cells: 10 Years On. Cell, 174(5), 1054–1066. 10.1016/j.cell.2018.07.017

Wang, H., Chen, H., Fu, Y., Liu, M., Zhang, J., Han, S., Tian, Y., Hou, H., & Hu, Q. (2022). Effects of Smoking on Inflammatory-Related Cytokine Levels in Human Serum. *Molecules (Basel*, Switzerland), 27(12), 3715. 10.3390/molecules27123715

Watson, A. M., Benton, A. S., Rose, M. C., & Freishtat, R. J. (2010). Cigarette smoke alters tissue inhibitor of metalloproteinase 1 and matrix metalloproteinase 9 levels in the basolateral secretions of human asthmatic bronchial epithelium in vitro. Journal of Investigative Medicine: The Official Publication of the American Federation for Clinical Research, 58(5), 725–729. 10.231/JIM.0b013e3181db874e

Worthington, A. K., Cool, T., Poscablo, D. M., Hussaini, A., Beaudin, A. E., & Forsberg, E. C. (2022). IL7Rα, but not Flk2, is required for hematopoietic stem cell reconstitution of tissue-resident lymphoid cells. Development (Cambridge, England), 149(8), dev200139. 10.1242/dev.200139

Zalokar, J. B., Richard, J. L., & Claude, J. R. (1981). Leukocyte Count, Smoking, and Myocardial Infarction. New England Journal of Medicine, 304(8), 465–468. 10.1056/NEJM198102193040806

Zhang, H., Rodriguez, S., Wang, L., Wang, S., Serezani, H., Kapur, R., Cardoso, A. A., & Carlesso, N. (2016). Sepsis Induces Hematopoietic Stem Cell Exhaustion and Myelosuppression through Distinct Contributions of TRIF and MYD88. Stem Cell Reports, 6(6), 940–956. 10.1016/j.stemcr.2016.05.002

